# Rapid deep widefield neuron finder driven by virtual calcium imaging data

**DOI:** 10.1101/2022.01.25.474600

**Authors:** Yuanlong Zhang, Guoxun Zhang, Xiaofei Han, Jiamin Wu, Ziwei Li, Xinyang Li, Guihua Xiao, Hao Xie, Lu Fang, Qionghai Dai

## Abstract

Widefield microscope provides optical access to multi-millimeter fields of view and thousands of neurons in mammalian brains at video rate. However, calcium imaging at cellular resolution has been mostly contaminated by tissue scattering and background signals, making neuronal activities extraction challenging and time-consuming. Here we present a deep widefield neuron finder (DeepWonder), which is fueled by simulated calcium recordings but effectively works on experimental data with an order of magnitude faster speed and improved inference accuracy than traditional approaches. The efficient DeepWonder accomplished fifty-fold signal-to-background ratio enhancement in processing terabytes-scale cortex-wide recording, with over 14000 neurons extracted in 17 hours in workstation-grade computing resources compared to nearly week-long processing time with previous methods. DeepWonder circumvented the numerous computational resources and could serve as a guideline to massive data processing in widefield neuronal imaging.

---

Development of optical microscopy^1-3^ and genetically encoded calcium indicators (GECIs)^4^ help researchers study the brain functionality in various behavior tasks, which further inspire the evolution of artificial intelligence (AI)^5^. For neuronal acquisitions within the scattering brain, a fundamental limitation is the trade-off between serial and parallel acquisition schemes^6^. Serial acquisition approaches such as two-photon laser-scanning microscopy (TPLSM) provide optical sectioning and robustness to scattering^7^, but are restricted in low temporal resolution and small field-of-view (FOV). Although recently developed multiplexing methods largely increase the TPLSM frame rate, the high power dosage in the animal brain^8^ could induce heat problems and irreversible damages. In the dimension of spatial accessibility, sophisticated optical design has pushed the TPLSM FOV to ∼5 mm in diameter^9^, but typically requires temporally sub-sampling of calcium dynamics for a cortex-wide region-of-interest (ROI). On the other hand, parallel schemes such as widefield microscope use cameras to detect widely distributed signals^6,10-12^, allowing video-rate acquisition over multi-millimeter scaled ROIs with single-cell resolution^13^. With optimized optical setup and computational tools, recently reported one-photon microscope has achieved 10 × 8 mm^2^ FOV in 0.8 µm resolution, which can cover tens of mammalian brain regions^14^ and hold great potential to record millions of neurons simultaneously^10^. The simplicity and high throughput of widefield microscope make it also popular in head-mounted microscope^15,16^, which is a compact and low-cost tool for studying social and behavior related activities. However, as the widefield microscope illuminates and detects the whole volume of the sample, neurons away from the focal plane contribute ambiguous background signals massively^17^. Light scattering in the opaque tissue further mixes fluorescence signals originating from the focal plane and confuses information about neuron locations. The inevitable background contaminations and scattering pose a big challenge to neuronal activity inference in the parallel acquisition scheme.

Computational approaches have been developed to separate neuronal signals from background contaminations in widefield microscope. The mostly-used constrained nonnegative matrix factorization (CNMF-E) approach models the strong background signals with prior knowledge of the spatial-temporal signal properties^18^. However, refining the background model for widefield imaging concomitantly requires sophisticated parameter tuning and huge computation consumption, preventing it from applications to cortex-scale neuron processing^19^. Online processing with a lightweight version of the algorithm partially alleviates the speed problem, but at the expense of performance downgrade^20^. Other methods^19,21,22^ without explicit modeling of the fluctuated background could achieve higher processing speed, but commonly face the risks of residual background contaminations^20^. Thus, it is still challenging to decipher widefield calcium recordings in scattering mammalian brains with both high speed and good performance using traditional computational methods.

The rapidly developing neural networks have achieved breakthroughs in neuronal image processing such as image enhancement^23^, neuronal segmentation^24,25^, and spike inference^26^. Recent research has demonstrated that, with proper training, deep learning enhanced neuronal activity inference can achieve an order of magnitude faster speed with no compromise of performance degradation^24^. This forecasts an opportunity of using neural networks processing functional data with downscaled time consumption. Yet in practice, seldom works have been reported that leverage the powerful deep learning to remove the background in widefield neuronal recordings, given the lack of paired widefield data and background-free captures. Methods that convert traditional background models into trainable convolutional filters alleviate the requirement of paired data, but need per-sample retraining and compromise the performance compared to traditional neuron extraction methods^20^.

Here we propose a fast and efficient neuronal extraction and demixing technique for the widefield microscope with nearly an order of magnitude faster speed and improved accuracy and precision through deep learning enhancement. Regarding the lack of training data, our method leverages a vivid simulation of the brain tissue^27^ to generate optical system-specific paired virtual recordings that with and without background. A neural network thus can be trained to separate neuronal signals from the scattered background (Methods, Fig. 1a, Supplementary Fig. 1, and Supplementary Video 1). A lightweight convolutional neural network (CNN) is then applied to quickly segment contamination-removed neurons to retrieve spatial footprints and temporal signals (Fig. 1b, Supplementary Fig. 2). We define the deep learning-inspired widefield neuron finder as DeepWonder. Demonstrated with both simulation and experiment data, we verified a nearly tenfold processing speed improvement with DeepWonder compared to the widely used CNMF-E algorithm. We further validated the accuracy of background removal and segmentation using DeepWonder by comparing with the TPLSM recordings of diverse animals *in vivo* on a hybrid system with simultaneous widefield and TPLSM recordings. We deploy DeepWonder on recently developed widefield calcium recording systems, including terabyte-scale RUSH system covering over 14,000 neurons, large FOV macroscope^13^, and head-mounted miniscope on freely-moving animals^16^. Finally, we package DeepWonder into a python package and distribute it in an online platform to make our method easy to access and convenient to use, for promoting interdisciplinary and reproducible researches.

**Fig. 1.**
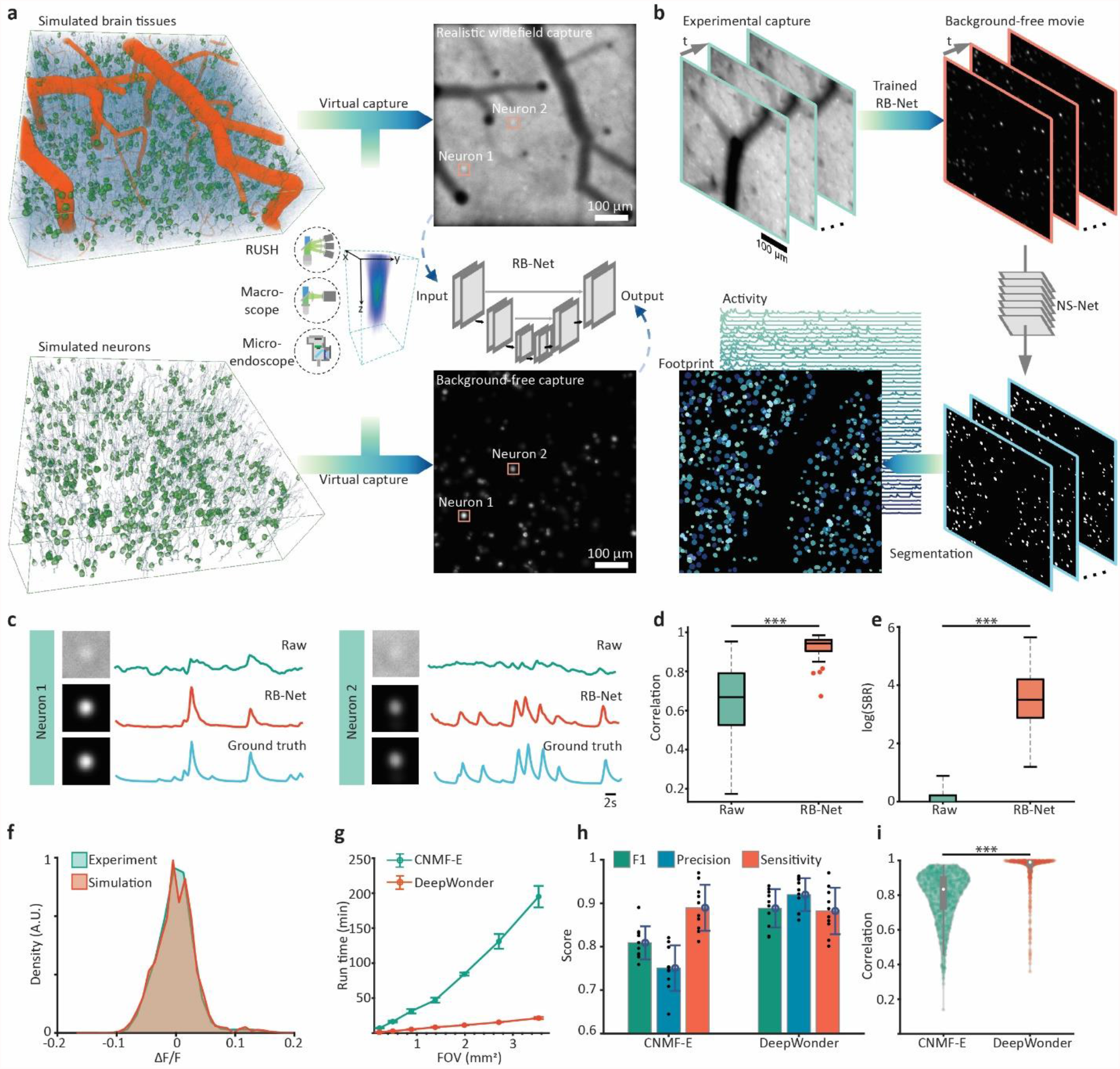
Principle of deep widefield calcium finder (DeepWonder). **a**. Training stage of removing background network (RB-Net) in DeepWonder. Based on specific widefield microscope parameters (numerical aperture, objective focal length, and magnifications) and imaging parameters (wavelength, imaging depths, and imaging power), we firstly use the proposed realistic widefield simulator to generate virtual captures with high similarity with experimental captures as inputs. Meanwhile, we can generate the capture with the same neuron distributions but without background contaminations as labels. We feed both inputs and labels to train the removing background network (RB-Net) such that it can restore background-free neuronal images from background-contaminated images. **b**. DeepWonder works on new recordings. After the network is trained, it can be used to remove the background of experimental captures. We further apply a neuron segmentation network (NS-Net) to segment neurons and extract neuronal signals from the background-removed movies (Supplementary Fig. 1, 2). **c**. Restoration of indiscernible calcium transients from backgrounds (green) by DeepWonder (red). Traces without background contamination (blue) serve as ground truth for comparison. **d**. Background removal of DeepWonder significantly increases the neuronal signal correlations with ground truth movie (p<0.001, two-sided Wilcoxon signed-rank test, n = 901; Methods). **e**. Background removal of DeepWonder significantly increases the signal-to-background (SBR) ratio (p<0.001, two-sided Wilcoxon signed-rank test, n = 901; Methods). **f**. distributions of simulation datasets (red) and experimental datasets (green). **g**. Increasing the FOV size leads both runtimes of CNMF-E and our approach quadratically increasing, but our approach is ∼10 times faster than CNMF-E in processing recordings in 2.0 × 1.7 mm^2^ FOV. Left panels show examples of MIP images of RUSH recordings in 0.22, 1.38, and 3.54 mm^2^. All movies are with constant 2000 frames. The plot show mean ± s.d. runtime, averaged over n = 5 different datasets (Supplementary Fig. 11). **h**. F1, precision, and sensitivity scores of segmentation by CNMF-E are 0.81± 0.03, 0.75 ± 0.05, and 0.89 ± 0.04, respectively. F1, precision, and sensitivity scores of segmentation by DeepWonder are 0.90 ± 0.03, 0.92 ± 0.03, and 0.88 ± 0.04, respectively. Statistical scores are shown in mean ± std across n=10 videos. DeepWonder is significantly better than CNMF-E in F1 score and precision (p=0.002, two-sided Wilcoxon signed-rank test). **i**. Correlation with ground truth of DeepWonder (red, 0.96 ±0.09, mean ± std across n=10 videos) and CNMFE (green, 0.79 ±0.14, mean ± std across n=10 videos). DeepWonder is significantly better than CNMF-E in correlations (p<0.002, two-sided Wilcoxon signed-rank test, n=1428).

The achievable neuron detection sensitivity and signal extraction quality in widefield microscope are ultimately limited by background contaminations, which mixed with crosstalk among neurons, neuropils, and background fluorescence from out-of-focus depths. In DeepWonder, we wipe out those contaminations by establishing a neural network that maps background contaminated images into background-free data (Fig. 1a). We generate realistic synthetic widefield calcium imaging data by adapting full modeling of vessels, neurons, and background dendrites and axons with a specific widefield microscope model, yielding virtual recordings with highly realistic pixel distribution,Δ*F*/*F* distribution, and spatial frequency distribution (Methods, Supplementary Fig. 3-5). In counterparts, background-free recordings are generated by only reserving fluorescent neuron and non-fluorescent vessels in the tissue. Paired virtual recordings are thus generated and fed to the proposed removing background network (RB-Net, Supplementary Fig. 1), to learn the mapping between domains of contaminated captures in real states and domains of background-free but never-existed captures. Trained RB-Net in DeepWonder learns interpretable features (Supplementary Fig. 6) and outputs high contrast images and vivid neuronal activities without contaminations (Fig. 1c). Compared to raw data, DeepWonder significantly enhances correlation scores to the ground truth signals (Fig. 1d, p<0.001, two-sided Wilcoxon signed-rank test, n = 901 neurons) and signal-to-background ratios (SBR) in test datasets that never been seen by the network (Fig. 1e, p<0.001, two-sided Wilcoxon signed-rank test, n = 901 neurons). Compared to other state-of-the-art background removal methods^19,20^, our RB-Net achieves superior performances in terms of SBR (Supplementary Fig. 8f), correlation score (Supplementary Fig. 8g), and neuron finding scores on the same dataset (Supplementary Fig. 8h, Supplementary Note 1), while spending almost 7-fold shorter time in removing background (Supplementary Fig. 8i). Guaranteed by the high similarity between virtual generations and real recordings (Fig. 1f), the RB-Net driven by virtual data in DeepWonder can be effectively applied to remove backgrounds of real recordings (Fig. 1b, Supplementary Fig. 7, Supplementary Video 1). We illustrate an SBR improvement in real recordings reaches over fifty times compared to raw data across 1543 neurons by RB-Net (Supplementary Fig. 7).

After separating neuronal signals from background contaminations, we then propose a neuron segmentation network (NS-Net) which efficiently segments neurons from background removed data (Methods, Fig. 1b). The NS-Net starts with a lightweight CNN that segments neurons from RB-Net output at a high speed (Method, Supplementary Fig. 1b). Roughly isolated neurons are further semantically segmented based on their spatio-temporal connectivity and turn to mostly exclusive segmentations. The temporal activities of those individual neurons can be directly read out since there is no inter-neuron crosstalk (Supplementary Fig. 2a). Neurons that are tiled and overlapped will be further demixed by a local nonnative matrix factorization (NMF)^28^ algorithm to eliminate activities crosstalk (Methods, Supplementary Fig. 2b). NS-Net reliably demixes neurons that are as close as 0.3 of the neuron diameter, yielding a temporal similarity over 0.9 and a spatial similarity over 0.85 (Supplementary Fig. 9). Our proposed NS-Net beats state-of-the-art two-photon segmentation technique CNMF and SUNS with the highest sensitivity and F1 score (0.933 and 0.917, respectively) in the background removed dataset (Supplementary Fig. 10e). The processing speed of NS-Net is 5 times faster than CNMF, and is comparable with SUNS (Supplementary Fig. 10f).

So far, by combining the optimized RB-Net and NS-Net into one framework, our DeepWonder eventually achieves a processing speed improvement of nearly ten folds (Fig. 1g, Supplementary Fig. 11b) compared to widely-used CNMF-E technique in widefield calcium imaging analysis. DeepWonder additionally brings out segmentation and activity inference accuracy improvement, represented as 11.1% promotion in F1 scores (Fig. 1h) and 21.5% promotion in temporal correlation scores (Fig. 1i). The proposed RB-Net in DeepWonder circumvents the time-consuming background modeling process in CNMF-E and achieves background elimination through a single-shot workflow, where the processing speed is only affected by the scale of datasets. Even higher speed acceleration is observed when cell density and cell number are higher, typically reaching nearly 20 times improvement when the neuron density reaches 5000 cells/mm^2^ (Supplementary Fig. 11a). Illustrated by the processing of calcium recordings of over 10,000 frames at 10 Hz, CNMF-E takes over two hours on average, while DeepWonder takes only 11 minutes (Supplementary Fig. 11c). DeepWonder is also robust to noise and reaches 0.60 F1 scores and 0.77 temporal correlations in a condition with an ultralow post-objective excitation power of 0.3mW/mm^2^, which is 9 folds and 1.6 folds higher than CNMF-E, respectively (Supplementary Fig. 12). In moderately low excitation power situations (0.7 mW/mm^2^), DeepWonder still outperforms CNMF-E in accuracy with F1 scores of 0.82 relative to 0.66 with CNMF-E.

To evaluate the inference accuracy of our proposed DeepWonder driven by simulated datasets, we next compare its performance with that of a standard two-photon microscopy. We build a hybrid microscopy device capable of both two-photon and widefield detection modalities. We sequentially switched the co-axis aligned two-photon and one-photon lightpath by timing control of a gated electrical optical modulator (EOM), LED excitation, and photon-sensitive photomultiplier (PMT) shutter in 30Hz (Fig. 2a). The shutter was used to protect the sensitive PMT when strong widefield fluorescence was excited (Supplementary Fig. 13). We reduced the two-photon excitation NA to 0.27 (Supplementary Fig. 14) such that the same neuron population could be detected by both the widefield and two-photon modalities. With image registration, we achieved 15Hz widefield neuronal recordings and paired 15 Hz two-photon recordings served as functional ground truth (Supplementary Note 2). We found the RB-Net in DeepWonder effectively mapped background-overwhelmed widefield data into sharp ones similar to the two-photon recordings in both spatial profile (Fig. 2b) and temporal activities (Fig. 2b and 2c). The correlation scores of DeepWonder output with two-photon signals reached 0.87±0.10, along with significantly reduced background contaminations (Fig. 2d; p<0.001, two-sided Wilcoxon signed-rank test, n=27; Supplementary Fig. 15, 16). With DeepWonder, we detected 47 neurons with 44 of them highly matched with active neurons from two-photon data, leading to F1 score of 0.91 compared to 0.73 by CNMF-E (Fig. 2e). By analyzing on 5 mice and 20 datasets, DeepWonder achieved over 0.8 median correlation scores in each of the datasets (Fig. 2f), and 0.88 ± 0.05 (mean ± std) precision scores (Fig. 2g) across all datasets, indicating that DeepWonder provides accurate neuronal segmentation and activity inference in mouse recordings. Further compared to widely used CNMF-E, DeepWonder stands out with both higher accuracy (F1 = 0.88 ± 0.04 of DeepWonder compared to F1 = 0.73 ± 0.13 of CNMF-E, n = 20; Fig. 2h) and higher signal correlations with two-photon ground truth (Fig. 2i), demonstrating advantages in both speed and performance.

**Fig. 2.**
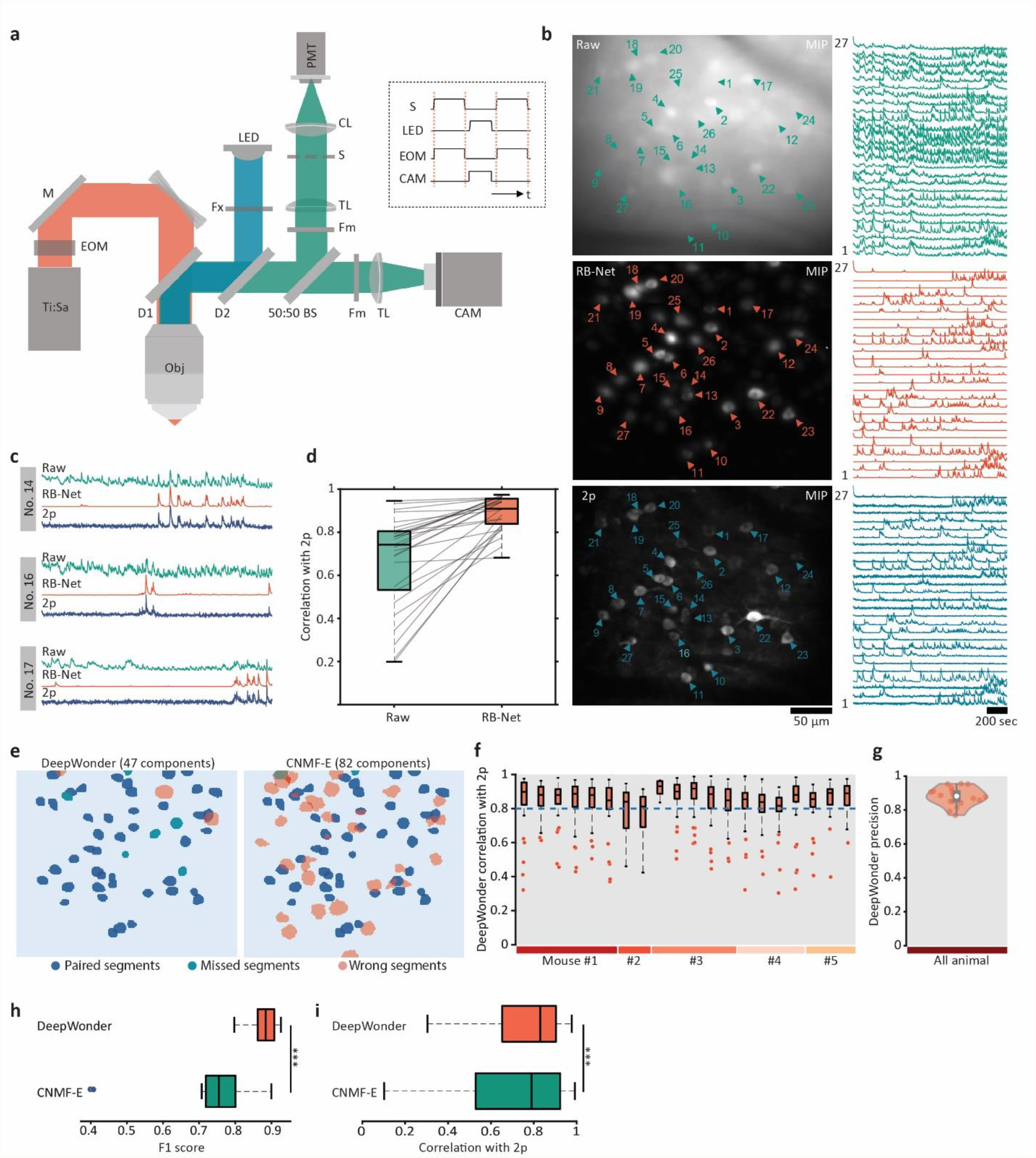
DeepWonder achieves accurate neuron segmentation and activity inference validated by two-photon (2p) microscope. **a**. The hybrid 1p–2p microscope setup. LED, light-emitting diode light source; Ti:Sa, titanium:sapphire laser; EOM, electro-optical modulator; M, mirror; DM, dichroic mirror; BS, beam splitter; Fm, emission filter; Fx, excitation filter; CL, collection lens; TL, tube lens; S, triggerable shutter; CAM, sCMOS (scientific complementary metal-oxide semiconductor) camera; PMT, photomultiplier tube; Obj, objective. Right box: control signals of the shutter, LED, EOM, and camera exposure, where high-level signals are activated and low-level signals are deactivated. **b**. Maximum intensity projection (MIP) of widefield (top), RB-Net processed widefield movie (middle), and two-photon movie (bottom). Triangles mark neurons and the corresponding temporal activities are plotted on the right side. **c**. Zoom-in plots of temporal activities of neuron No. 14, 16, and 17 in the widefield raw movie (green), RB-Net de-background movie (red), and 2p movie (blue.) **d**. Temporal correlations of 27 picked neurons with 2p by RB-Net are significantly increased compared to the raw movie (p<0.001, two-sided Wilcoxon signed-rank test). **e**. DeepWonder (left) and CNMF-E segmentation results (right). Blue masks represent corrected segmentations, green masks represented missed segmentations by current methods, and pink masks represent false segmentations by current methods. The precision, sensitivity, and F1 score of DeepWonder are 0.94, 0.88, and 0.91, while for CNMF-E are 0.56, 0.96, and 0.73. **f**. Correlations of DeepWonder neuron activities with 2p across 5 animals and 20 datasets. **g**. Precision score of DeepWonder segmented neurons with 2p dataset as the reference across 5 animals and 20 datasets reaches 0.88 ± 0.05 (mean ± std). **h**.F1 score of DeepWonder (red, 0.88 ± 0.04, mean ± std) and CNMF-E (blue, 0.73 ± 0.13, mean ± std) across all datasets. DeepWonder significantly outperforms CNMF-E in F1 score (p<0.001, two-sided Wilcoxon signed-rank test). **i**. Temporal correlation of DeepWonder (red, 0.83 ± 0.14, median ± median absolute deviation) and CNMF-E (0.79 ± 0.22, median ± median absolute deviation) with 2p across all datasets (n = 1570 neurons). DeepWonder significantly outperforms CNMF-E in temporal correlations with 2p (p<0.001, two-sided Wilcoxon signed-rank test).

The enhanced computational efficiency of DeepWonder enables us to proceed cortex-wide neuronal recording within acceptable time-spent. Here, we demonstrate the data processing ability of DeepWonder on the terabytes-scale RUSH system^14^. The RUSH system is consisted of tens of sCMOS cameras, with totally 14800 × 15200 pixels across 10 × 8 mm^2^ FOV in 0.8 µm resolution at video rate, fertilizing population-scale neuron connection inference (Methods). We virtually generated lifelike neuron recordings based on optical parameters of the RUSH system (Supplementary Fig. 3), and trained DeepWonder for the data modality by the RUSH system. With DeepWonder, neurons that are overwhelmed by highly fluctuated backgrounds are clearly unveiled (Fig. 3a; Supplementary Video 3), and high-contrast calcium transients are uncovered (Fig. 3b) thanks to effective background suppression (Supplementary Fig. 17a, b, Supplementary Fig. 7). The high data throughput by the RUSH system yields over 1 terabyte of data in a 13.5 minutes imaging session at 10 Hz (Methods). Processing such a huge dataset with the popular CNMF-E technique takes over five days to fully demix neuron activities (132.4 hours in total, without counting loading time; Fig. 3c). In contrast, engined with highly accelerated DeepWonder, data of the same scale can be analyzed and inferred within 17 hours. Up to 14,226 neurons across 9 cortical areas were found with clear activities (Fig. 3d), showing great potential for interrogating behavior-related neuron population response within multiple cortical regions. When the awake mouse was anesthetized in the fifth minute with 2% isoflurane^29^, we observed neurons gradually became inactive across different cortical regions with different dynamics (Fig. 3d, Supplementary Fig. 20). We further manually annotated neurons in a small FOV (∼450 µm x 450 µm), and found DeepWonder achieved superior neuron segmentations than CNMF-E (Fig. 3e). The segmented neurons by DeepWonder are more concentrated in a round shape compared to those segmented with CNMF-E (Fig. 3f, Supplementary Fig. 17i), and the extracted calcium activities are with a higher signal-to-noise ratio (Fig. 3g, Supplementary Fig. 17h). DeepWonder achieves 0.87 ± 0.10 F1 scores in finding valid neurons compared to 0.74 ± 0.06 (mean ± std, n = 5; Fig. 3h, Supplementary Fig. 17e) by CNMF-E, indicating improved neuron segmentation accuracy in addition to the largely accelerated speed.

**Fig. 3.**
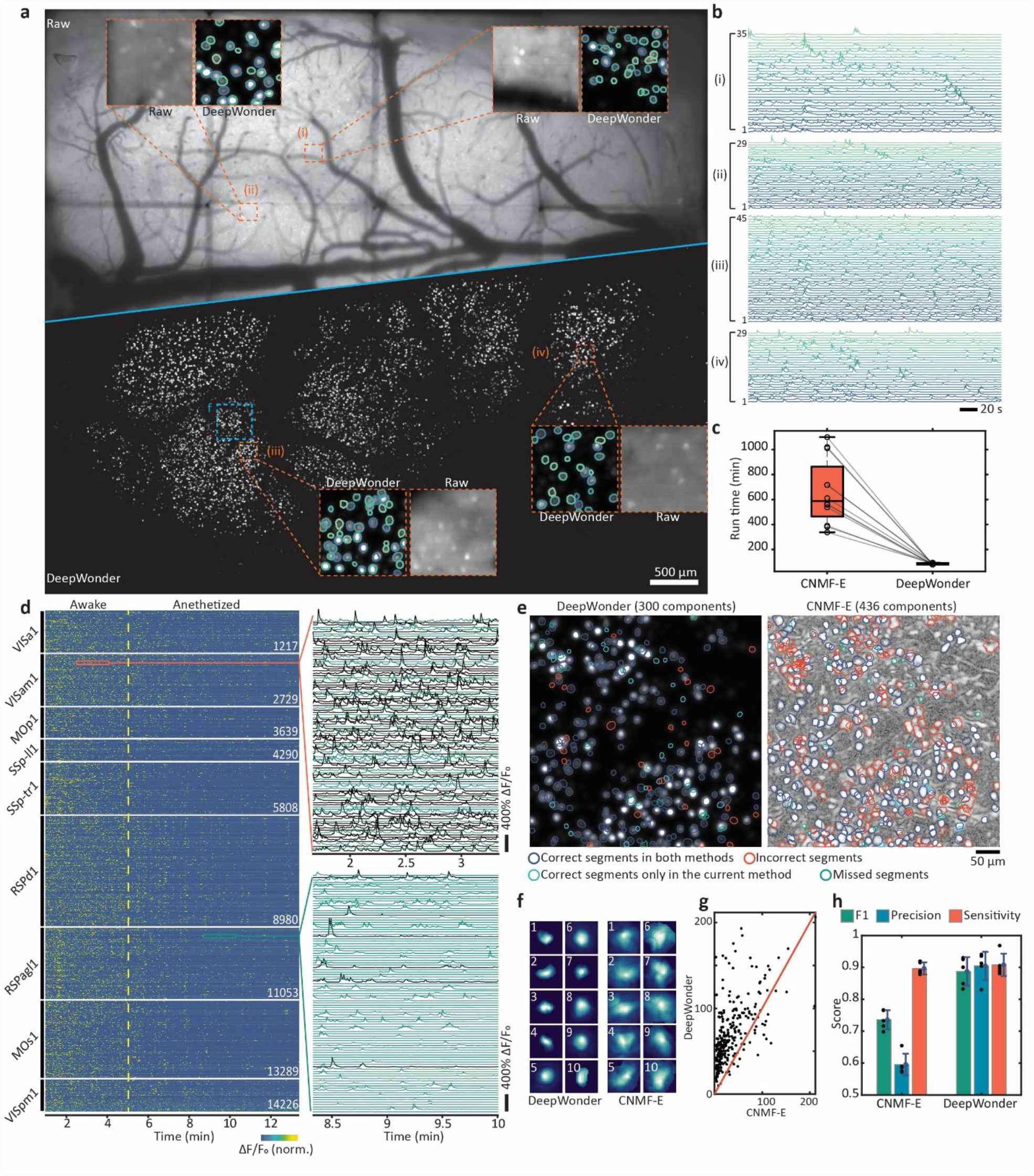
DeepWonder realizes high-speed processing widefield neuronal recordings in terabytes scale. **a**. Maximum intensity projection (MIP) of raw RUSH video (top) and background removed movie by DeepWonder (bottom). Orange dashed boxes mark zoom-in four areas in the cortex, with DeepWonder segmentations overlaid. **b**. Inferred calcium activities from four different areas marked by the dashed box with neuron number labeled. **c**. Runtime comparisons between CNMF-E and DeepWonder across 12 valid FOVs in RUSH over 8000 frames recordings. Each dot shows the processing time for each FOV. **d**. Temporal activity rendering of 14226 neurons inferred by DeepWonder in a 13.5 minutes recording. Two zoom-in panels show example traces (each with 100 traces). The dashed yellow line indicates the dosage of 2% isoflurane for anesthesia at 5 minutes after the secession start. **e**. The contour plot of all neurons detected by DeepWonder (left) and CNMF-E (right) superimposed on the standard deviation (STD) of background removed images and correlation image, respectively (Methods), respectively. Compare to manual segmentations, deep blue circles mark correct segments in both methods, red circles mark incorrect segments in each of the methods, green circles marks missed segments, and shallow blue circles mark correct segments that are only in the current method. **f**. Spatial components of 10 example neurons detected by both DeepWonder (left) and CNMF-E (right). **g**. The signal-to-noise ratio (SNRs) of all neurons detected by DeepWonder (vertical axis) and CNMF-E (horizontal axis) in **e**. **h**. F1, precision, and sensitivity scores of segmentation in **e** by CNMF-E are 0.74 ± 0.06, 0.58 ± 0.07, and 0.88 ± 0.04, respectively. F1, precision, and sensitivity scores of segmentation by DeepWonder are 0.87 ± 0.10, 0.91 ± 0.09, and 0.88 ± 0.07, respectively. Statistical scores are shown in mean ± std across n=5 samples.

The DeepWonder is designed to be a general technique that can be compatible with widely used widefield calcium imaging platforms including whole-brain macroscopy and miniaturized microscopy for freely-moving animals. In a macroscope with a photographic lens as the objective^13^, neurons are largely under sampled by ∼5×5 pixels laterally to redeem a multi-millimeter FOV. We virtually generated vivid neuron recordings based on magnification, numerical aperture, and other optical parameters of the macroscope system (Methods, Supplementary Fig. 4) to train DeepWonder. We found DeepWonder significantly reduced fluctuated backgrounds and highlighted neurons efficiently (Supplementary Fig. 18a, b, Supplementary Video 4). DeepWonder achieved 0.88 F1 scores compared to 0.81 by CNMF-E with manual labeling as ground truth (Supplementary Fig. 18c, d). Neurons found by DeepWonder only showed high-contrast calcium dynamics and compact shapes (Supplementary Fig. 18f). Head-mounted microendoscope is another widely used one-photon functional imaging technique that suffers from highly fluctuated backgrounds. With DeepWonder trained by virtual data in the microendoscope modality (Methods, Supplementary Fig.5), background contaminations were largely reduced (Supplementary Fig. 19a, b, Supplementary Video 5). Compared to manual inspection, DeepWonder achieved 0.91 F1 scores compared to 0.84 by CNMF-E with improved SNRs (Supplementary Fig. 19c-e). The effectiveness of DeepWonder across multiple platforms states the great potential of accelerating analysis of various widefield neuronal recordings.

In summary, DeepWonder accomplishes widefield functional inference with an order of magnitude faster speed, 11.1% improved F1 scores, and 21.5% improved temporal correlation scores. This performance is enabled by key advances in our unique solution for handling background contamination data: firstly, our proposed virtual widefield data generators can adapt to various optical microscopes and produce virtual recordings with high similarity in both pixel distributions and functional distributions. Secondly, paired virtual recordings with and without background contaminations are used to train a time-aware removing background network (RB-Net) that effectively peels background signals in experimental recordings. Thirdly, our neuron segmentation network (NS-Net) effectively processes both non-overlapped and overlapped components in a background-removed movie with 5 times faster speed than state-of-the-art technique. We validate the accuracy of DeepWonder in a customized simultaneous one-photon and two-photon observation system across multiple animals, and infer terabytes neuronal recordings within 17 hours via workstation-grade computing resources compared to the week-long processing time with traditional methods. To maximize its accessibility, we have open-sourced both DeepWonder and the widefield data generator to promote interdisciplinary researches.

It is worth noting that paired widefield and two-photon functional data can be acquired by the proposed hybrid system, based on which an end-to-end neural network can be trained to directly map widefield frames to background-free frames. However, we declare that DeepWonder driven by virtual calcium recordings outperforms the end-to-end model for the following reasons. Firstly, inactive neurons with low intensity that are invisible in widefield captures are visible in two-photon captures, posing difficulties to the network. On the contrary, synthetic datasets based on vivid tissue simulation and real imaging model cleanse inactive parts from labels and let the network focus on active neuron generation. Secondly, pixel-level alignment of widefield and background-free captures is critical for network training, which is readily guaranteed using synthetic data but difficult to achieve using two-photon data as labels. Thirdly, the cross-modality network training approach requires a hybrid imaging system as described in the article that is complicated to build, cost unfriendly, and even inapplicable (e.g., head-mounted microscope), whereas the DeepWonder can be effectively applied to any widefield system without extra efforts.

By entirely avoiding the hand-crafted model of complex neuronal background that in previous widefield functional inference algorithm^18^, the DeepWonder concept is also positioned to analyze the signals with higher temporal bandwidth (∼500 Hz) offered by genetically encoded voltage^30^ or functional imaging with other indicators^31^, simply by changing formulation of virtual recordings. Utilizing generative adversarial networks (GANs) holds potentials to further improve performances of DeepWonder^32^. With a further intaking model of widefield imaging in over hundreds of microns cortical depth^33^, DeepWonder scheme can also be expected to increase the performance of neuron extraction and activity inference in deeper cortical layers, potentially overcoming the current limitations of widefield imaging in superficial regions of the mammalian cortex. On the other hand, by reinforcing DeepWonder with volumetric imaging models such as light-field microscope^34^ and multifocus microscope^35^, DeepWonder can be further extended to infer volumetric neuronal activities at high speed. We anticipate the proposed method lowers the barrier of processing neuronal data by high-throughput and large-scale widefield captures, and promote whole brain and million level neuronal recordings.

## DATA AVAILABLITY

We have mounted our data in Google Colab, which is a free Jupyter notebook environment that requires no setup and runs entirely in the cloud. A demo script with full processing of DeepWonder on several demo datasets (including NAOMi1p virtual datasets, cropped RUSH datasets, and two-photon validation datasets) is available through Colab via https://colab.research.google.com/drive/15TvsyEYgE1iGpaNWkq3flXOw52I51mVa. All other data of this study are available from the corresponding author on request.

## CODE AVAILABLITY

Our DeepWonder with realistic widefield imaging simulators can be found at https://github.com/yuanlong-o/Deep_widefield_cal_inferece. An archived version of DeepWonder packages is available through https://pypi.org/project/DWonder.

## ACKNOWLEDGMENTS

We thank Yiliang Zhou and Zhifeng Zhao for helping in setting up the joint two-photon and one-photon microscope. This work was supported by the National Natural Science Foundation of China (No. 62088102), Ministry of Science and Technology of the People’s Republic of China (No. 2020AA0105500).

## AUTHOR CONTRIBUTIONS

Y.Z. designed and conceptualized the DeepWonder pipeline, performed two-photon validation experiments, and wrote the manuscript. G.X. implemented the DeepWonder pipeline, performed simulations, analyzed data, and wrote the manuscript. X.H. contributed to macroscope imaging, two-photon validation experiments, and analyzed data. J.W. and X.L. provided critical support on system setup and imaging procedure. Z.L. contributed to the final version of the manuscript. G.H. and H.X. performed cranial window surgeries, viral injections, RUSH imaging, and contributed to the manuscript. L.F. and Q.D. conceived and led the project and wrote the manuscript.

## METHODS

### One-photon and two-photon joint validation

To valid our algorithms in achieving correct neuronal activities, we built a joint two-photon and widefield detection system. The system was based on a standard two-photon laser scanning microscope (TPLSM), while we further added a 470 nm-centered widefield illumination path and a camera detection path in the system. The schematic of the custom-built two-photon microscope is shown in Supplementary Fig. 13. A titanium-sapphire laser system (MaiTai HP, Spectra-Physics) was served as the two-photon excitation source (920 nm central wavelength, pulse width <100 fs, 80 MHz repetition rate). A half-wave plate (AQWP10M-980, Thorlabs) and an electro-optic modulator (350-80LA-02, Conoptics) were used to modulate the excitation power. A 4f system (AC508-200-B and AC508-400-B, Thorlabs) with a 2x magnification was used to expand the laser beam to a resonant scanner (8315K/CRS8K, Cambridge Technology). The scanned beam went through a scan lens (SL50-2P2, Thorlabs) and a tube lens (TTL200MP, Thorlabs) and formed a tight focus through a high numerical aperture (NA) water immersion objective (25x/1.05 NA, XLPLN25XWMP2, Olympus). A high-precision piezo actuator (P-725, Physik Instrumente) drove the objective for fast axial scanning. To match the two-photon excitation range with the widefield detection range, we reduced the beam size at the back aperture of the objective with an iris. The effective excitation NA was about 0.27 in our imaging experiments, yielding ∼20 um axial range (Supplementary Fig. 14). A long-pass dichroic mirror (DMLP650L, Thorlabs) was used to separate fluorescence from femtosecond laser beam by reflecting the fluorescence signals and transmitting the infrared light.

For the widefield excitation path, a long pass dichroic (DMLP505L, Thorlabs) in the original detection path of TPLSM was used to send blue LED light (M470L4-C1 and MF475-35, Thorlabs) to the objective. To jointly record widefield excitation and two-photon excitation, a 50:50 (reflectance: transmission) non-polarizing plate beam splitter (BSW27, Thorlabs) was placed after widefield dichroic to separate fluorescent signals for PMT (PMT1001, Thorlabs) and camera (Zyla 4.2, Andor). A pair of fluorescence filters (MF525-39, Thorlabs; ET510/80M, Chroma) was configured in front of both the PMT and the camera to fully block both femtosecond laser and widefield excitation beam. The back aperture of the objective was optically conjugated to the detection surface of the PMT with a 4f system (TTL200-A and AC254-050-A, Thorlabs).

To avoid excitation crosstalk and protect PMT from high-flux widefield emission photons, we added a linear galvo that served as an optical shutter for the PMT detection path, which deflected fluorescent photons when LED was on (Supplementary Fig. 13a). We further configured the EOM to be blocked during widefield imaging. The LED (M470L4-C1) was in trigger mode with a typical rising and falling time less than 1 ms, with further reduced duration time to avoid PMT overexposure (Supplementary Fig. 13b).

### Realistic widefield capture generation

To synthesize a realistic cortical tissue and generate corresponding widefield capture, we referred to the Neural Anatomy and Optical Microscopy (NAOMi)^16^ package. Using NAOMi, a brain tissue volume was populated with multiple blood vessels, as well as with neuron somata, axons, and dendrites. Neurons and dendrites were assigned synthesized fluorescence activity that reflected their calcium dynamics. A tissue-specific point spread function (PSF) was generated by layer-to-layer Fresnel propagations from deep tissue to sensor.

While original NAOMi was used to simulate two-photon excitations, here we modified the original NAOMi pipeline such that it could faithfully simulate data acquisition of one-photon excitations, which was termed as NAOMi1p. We changed the excitation wavelength from the near infrared range into the visible range. In two-photon microscope, scattering-induced aberrations in excitation beam instead of emission beam affect the imaging quality due to the point-scanning manner. On the other hand, in widefield microscope, scattering-induced aberrations cause troubles in emission beam instead of excitation beam due to the planar collection from different camera pixels. We thus modified the optical PSF generation based on the propagation of the emission beam instead of the excitation beam through the tissue. We further replaced the two-photon absorption process with one-photon absorption process in a model of power density, fluorescent concentration, extinction coefficient, quantum yield, and fluorescent protein expression level^36^. The final simulated recordings have three contributors: fluorescence from active neurons, fluorescence from dendrites and axons in the background, fluorescence from out-of-focus backgrounds. The assembly of all three parts faithfully generates a virtual capture of widefield recordings, while using fluorescence from active neurons only generates a background-free label. Especially, for soma target indicators^37^ it is recommended to only let the soma fire. The above tools are summarized as the NAOMi1p toolbox and are open to all the community. To accommodate different imaging systems, NAOMi1p opens multiple parameters including the acquisition NA, camera pixel size, magnifications, illumination power, FOV, and indicator types for users to adjust. With NAOMi1p, we can faithfully generate virtual widefield recordings as well as their background-free counterpart. We then picked up neurons that were within the range of axial PSF diameter (Gaussian beam, 1/e^2^ size) and registered their positions and activities as ground truth for simulation comparisons among different analysis algorithms for widefield calcium recordings (Supplementary Note 1).

### Noise simulation

Imaging sensors (e.g. sCMOS, CMOS, and CCDs) have different quantum efficiency (QE) and noise response, which is also highly coupled with the expression level of calcium indicators in neurons. We thus simulated the NAOMi tissue with a range of noise to cover those situations. The number of fluorescence photons generated in a unit area of the samples is^36^

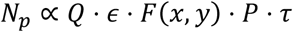

where *Q* is the quantum efficiency of fluorophores with an extinction coefficient *∈, F* (*x,y*) is the local fluorophore concentration,*P* is power density, and *τ* is the integration time of the camera. The signal of a camera can be further interpreted as^38^

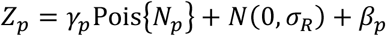

where *γ*_*p*_ is a multiplicative factor which is applied to the Poisson-distribution (Pois) as the camera gain.*β*_*p*_ is a bias during analog-to-digital (AD) conversion. *N*(0,*σ*_*R*_) is the Gaussion-distribted readout noise with zero mean and *σ*_*R*_ standard deviation. For a typical sCMOS, *γ*_*p*_ is ∼2.2, *β*_*p*_ is ∼100, and *σ*_*R*_ is ∼ 200^39^.

### Widefield imaging setups and recordings

#### RUSH recordings

In the real-time, ultra-large-scale, high-resolution (RUSH) system^14^, a 5×7 customized field lens array was mounted on a spherical surface for full correction of field curvature of 10 mm × 12 mm FOV. The customized objective provides 0.35 NA across the centimeter scale FOV, supporting submicron resolution observation. The pixel resolution of each camera in RUSH system is 2560 × 2160, yielding 6.3 gigabyte data per minute at 10 Hz. A mouse with a 7 mm cranial window takes 12 sub FOVs of RUSH, and 13.5 minutes recordings take over 1 terabyte of data (Fig. 4, Supplementary Fig. 20). To generate virtual recordings for DeepWonder training, we fed the following typical parameters to the data generator: system magnification 10, NA 0.35, pixel size 0.8 µm, frame rate 10Hz, and illumination power density 0.8mW/mm^2^. We checked the similarity of generated data with raw recordings (Supplementary Fig. 3).

#### Macroscope recordings

##### Macroscope recordings

We used a 50mm camera lens (Canon EF 50mm f/1.4 USM) as the objective lens and a 100mm camera lens (MINILTA AF 100mm f/2.8) as the tube lens to set up the widefield macroscope. The illumination was provided by a collimated blue LED (SOLIS-470C, Thorlabs) with an excitation filter (FESH0500, Thorlabs). The beam was focused by a lens (AC508-100-A, Thorlabs), reflected by a dichroic mirror (DMLP505L, Thorlabs), passed through the objective lens and excited the sample. The fluorescence was collected by the same objective lens and refocused on the sCMOS camera (Zyla 5.5, Andor) by the tube lens. An emission filter (MF525-39, Thorlabs) was placed before the camera to eliminate the excitation light. The FOV of the system was approximately 9.2 mm x 7.7 mm, and each pixel in the sCMOS corresponded to 3.6 µm on the image plane. To generate virtual recordings for DeepWonder training, we fed the following typical parameters to the data generator: system magnification 1.8, NA 0.3, pixel size 3.6 µm, frame rate 10Hz, and illumination power density 0.8mW/mm^2^. We checked the similarity of generated data with raw recordings (Supplementary Fig. 4).

#### Head-mounted microscope recordings

The data of head-mounted microscope recordings was released by Wang Lab at McGovern Institute in MIT, and downloaded from https://github.com/JinghaoLu/MIN1PIPE. The data was recorded with open-source UCLA miniscope^16^. To generate virtual recordings for DeepWonder training, we fed the following typical parameters to the data generator: system magnification 6, NA 0.46, pixel size 1 µm, frame rate 20Hz, and illumination power density 2.5 mW/mm^2^. We checked the similarity of generated data with raw recordings (Supplementary Fig. 5).

### Network architecture and training

#### Removing background network

The main structure of removing background network (RB-Net) is 3D Unet. The encoding path and decoding path consist of three convolutional blocks (Supplementary Fig. 1a). For accelerating removing background process, we added a “spatial to channel” down-sampling operator at the beginning of RB-Net for reshaping the input image of size W×H×C into W/2×H/2×4C (W for filter width, H for filter height, and C for filter channels; Supplementary Fig. 1c). We also introduced a “channel to spatial” up-sampling operator at the end of RB-Net for realigning pixels (Supplementary Fig. 1c). With these two operators, the pixel number of an input image processed by RB-Net can be increased by four times at almost the same GPU memory cost. We utilize a linear transformation of raw input images *x* for data augmentation as:

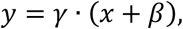

where *y* is input images for RB-Net, *γ* and *β* are random number (0.2 < *γ* < *β* < *max* (*x*)). Data augmentation is constructive to the generalization ability and transfer learning ability of RB-Net.

We synthesized 23 sets of background removed data by the NAOMi1p algorithm and randomly split them into 4000 paired patches for training RB-Net. The input raw videos were mean subtracted. It took 48 hours to train RB-Net for 30 epochs with a Geforce RTX 3080 GPU. The running speed for RB-Net is usually 40 ms per 750 × 750-pixel frame tested in an RTX 3080 GPU.

#### Neuron segmentation network

The main structure of neuron segmentation network (NS-Net) is 3D Unet, which has the similar structure with the RB-Net but with different channels (Supplementary Fig. 1b). On the other hand, because neuron segmentation in background-free data is simpler than removing background, we utilize the combination of a 1×1×3 filter and a 3×3×1 filter in NS-Net to replace two 3×3×3 filters for reducing network parameters and computational consumption.

The training data for NS-Net is directly generated from NAOMi1p generator, where neuron soma that are within the range of axial PSF diameter (Gaussian beam, 1/e^2^ size) is binarized as segmentation label. We simulated 45 sets of neuron segmentation data and randomly generated 4000 paired patches for training neuron segmentation network. We spent 8 hours training neuron segmentation network for 30 epochs with a Geforce RTX 1080TI GPU.

### Processing of widefield calcium data

Widefield calcium recordings are firstly sent to trained RB-Net to get a de-background clean movie, then the background-free movie is further sent to trained NS-Net for acquiring neuron candidate masks (Supplementary Fig. 2a). We group and merge candidates from all frames into connected regions to form unique segmentations. We then do the connectivity analysis for every candidate of the mask sequence spatio-temporally and extract every separated neuron to compose a neuron candidate list. Those spatially overlapped but temporally separated (e.g. neuron segments appear in different frames) will be registered as different candidates. With the neuron candidate list, we classify these neurons by neuron morphology metrics related to area and roundness 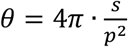, where *s* is the area of neuron, *p* is the perimeter of neuron. We abandon the neuron candidates that are smaller than the 25 µm^2^ threshold. Since the roundness *θ* is a good indicator to judge if the candidate consists of a single neuron or multiple neurons, we further classify neuron candidates whose roundness are higher than the standard roundness of a single neuron (typically *θ* =0.8) to form a “good” neuron list, and others into a “bad” neuron list (Supplementary Fig. 2b). The candidates in the “good” neuron list will be sent out for directly reading out temporal activities from the background-removed movie based on values of exclusive pixels (Supplementary Fig. 2a). For each candidate in the “bad” neuron group, the candidate will be initialized by greedy methods^18^ and then sent to local NMF for further demixing. If we mark the local area surrounding the candidate as 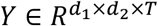, and the candidate is estimated to be consisted by *K* neurons, the local NMF model is then

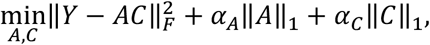

where 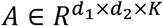 and *C ∈ R*^K×T^ represents the spatial and temporal footprints, respectively^18^. We solve the above optimization problem through HALS algorithms^28^. Finally, we merge neurons by clustering components with high temporal correlations and spatial overlap ratios.

### Mouse preparation and calcium imaging

All animal experiments were performed following institutional and ethical guidelines for animal welfare and have been approved by the Institutional Animal Care and Use Committee (IACUC) of Tsinghua University. Mice were housed in cages (24 °C, 50% humidity) in groups of 1–5 under a reverse light cycle. Both male and female mice were used without randomization or blinding. Adult transgenic mice (cross between Rasgrf2-2A-dCre mice (JAX 022864) and Ai148 (TIT2L-GC6f-ICL-tTA2)-D (JAX 030328)) at 8–12 postnatal weeks were anesthetized with 1.5% isoflurane, and craniotomy surgeries were conducted with a stereotaxic instrument (68018, RWD Life Science) under a bright-field binocular microscope (77001S, RWD Life Science). A custom-made coverslip fitting the shape of the cranial window was cemented to the skull. A biocompatible titanium headpost was then cemented to the skull for stabilization during imaging. The edge of the cranial window was enclosed with dental cement to hold the immersion water of the objective. After the surgery, trimethoprim (TMP) was injected into the mice intraperitoneally for inducing the expression of GCaMP6f in layer2/3 neurons (0.25mg per gram). To reduce potential inflammation, 5 mg per kg (body weight) of ketoprofen was injected subcutaneously. Each mouse was housed in a separate cage for 1–2 weeks of postoperative recovery.

Imaging experiments were carried out when the cranial window became clear and no inflammation occurred. Mice were first rapidly anesthetized with 3.0% isoflurane and then fixed onto a custom-made holder by the headpost. A precision 3-axis translation stage (M-VP-25XA-XYZL, Newport) carried the mice for a proper region of interest. For two-photon validation experiments, the correction ring of the 25x water immersion objective was adjusted to compensate for the coverslip thickness and eliminate spherical aberrations. The highest excitation power of two-photon microscope after the objective was under ∼100 mW to avoid head damage. During the imaging session, gaseous anesthesia was turned off and the mouse was kept awake. We performed single-plane imaging at approximately 80 μm below the pia mater. For widefield acquisitions, the excitation power density in the cranial window area was no more than 1.5 mW/mm^2^. Before running further analysis, we ran calcium movie registrations with open-source NormCorre algorithm^40^ to cancel motion artifacts. In cortex-wide brain imaging, we aligned the recorded brain area into Allen CCF atlas based on the recorded position of the cranial window by the stereotaxic instrument when applying brain surgery.

### Performance metrics

#### Correlation score

We used Pearson’s correlation coefficient as the temporal metric to monitor the similarity between inferred neuronal activities and ground truths. The ground truth activities were available for simulation data, while for joint one-photon and two-photon validation data, the ground truth activities were established by running CaImAn^41^ on two-photon dataset (Supplementary Note 2).

#### Neuron finding scores

It is necessary to establish ground truth segmentations for comparing the neuron finding scores. In simulation data, the ground truth segmentations were available. In joint one-photon and two-photon validation data, the ground truth segmentations were marked based on CaImAn processed two-photon data (Supplementary Note 2). In widefield experimental data, we manually labeled the neurons based on their positions and activities. We firstly calculated the correlation images of the raw recordings^41^, and worked over every structure that was different from the background which matched neuron size (typically 10∼15 µm in diameter). We rejected those candidates that with weak and noisy activities in the original movie. We outlined each cell of interest with the ROI manager in ImageJ, and imported the zipped ROIs into MATLAB as ground truths for comparisons with other methods.

After achieving segmentation ground truth, a customized script in MATLAB automatically judges segmentations by the following rules: a candidate is a correct segment (true positive, TP) if the minimal distance between this candidate with any ground truth segments is less than 8 µm, and the Intersect over Union (IoU) score between this candidate and any ground truth segments is larger than 0.2. Otherwise, the segmentation candidate will be rejected as a false positive (FP). Segments that are recognized by ground truth labeling but not recognized by the algorithm will be marked as false negatives (FN). The segmentation accuracy (F score,*F*1) is defined as

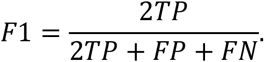

The segmentation precision score is defined as

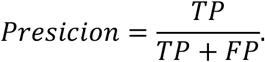

#### Signal-to-background ratio

We calculated the signal-to-background ratio (SBR) of a neuron by computing the maximum activity of the neuron area over the maximum activity of its neighboring area (Supplementary Fig. 7). The neuron area is defined by a circle with a radius of 10 µm with the center at the centroid of a segmentation. A neighboring area is defined by a ring with an inner radius of 10 µm and an outer radius of 20 µm at the same center of the corresponding neuron area, with masking out all other neuron areas.

#### Signal-to-noise ratio

We computed the signal-to-noise ratio (SNR) of inferred cellular traces to quantitatively compare the temporal inference quality^18^. We calculated the denoised trace of each inferred activity using OASIS^42^, and the SNR was computed through

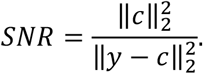

